# Non-invasive red-light optogenetic control of *Drosophila* cardiac function

**DOI:** 10.1101/2020.01.26.920132

**Authors:** Jing Men, Airong Li, Jason Jerwick, Zilong Li, Rudolph E. Tanzi, Chao Zhou

## Abstract

Drosophila is a powerful genetic model system for cardiovascular studies. Recently, optogenetic pacing tools have been developed to control *Drosophila* heart rhythm noninvasively with blue light, which has a limited penetration depth. Here we developed both a red-light sensitive opsin expressing *Drosophila* system and an integrated red-light stimulation and optical coherence microscopy (OCM) imaging system. We demonstrated noninvasive control of *Drosophila* cardiac rhythms, including simulated tachycardia in ReaChR-expressing flies and bradycardia and cardiac arrest in halorhodopsin (NpHR)-expressing flies at multiple developmental stages. By using red excitation light, we were able to pace flies at higher efficiency and with lower power than with equivalent blue light excitation systems. The recovery dynamics after red-light stimulation of NpHR flies were observed and quantified. The combination of red-light stimulation, OCM imaging, and transgenic *Drosophila* systems provides a promising and easily manipulated research platform for noninvasive cardiac optogenetic studies.

## Introduction

Cardiac optogenetics enables noninvasive optical stimulation of the heart with high spatial and temporal precision. It has been increasingly studied as an alternative to electrical stimulation methods for cardiac control since 2010.^1–5^ Arrenberg *et al.* first developed transgenic *Zebrafish* models with a genetically-encoded, optically-controlled pacemaker expressing channelrhodopsin-2 (ChR2) or halorhodopsin (NpHR) in the heart.^6^ Bruegmann *et al.* used ChR2-expressing cardiac tissue for light-induced stimulation of heart muscle *in vitro* and in mice with open chest preparation.^1^ Over the last few years, the field of cardiac optogenetics has experienced rapid growth.^4, 7–32^ Several groups, including ours, have been working on different frontiers in this field, and significant progress has been made. Important contributions were introduced by Nussinovitch *et al.*, who demonstrated optogenetic pacing and resynchronization therapies in surgically exposed rat hearts *in vivo*.^4^ Vogt *et al.* developed a novel gene transfer method to deliver ChR2 specifically to the adult mouse heart via a simple systemic injection of adeno-associated virus serotype 9 (AAV9).^21^ Our group developed transgenic fly models expressing ChR2 in the heart.^3^ For the first time, we demonstrated successful optogenetic pacing in fruit flies at different developmental stages, including larva, pupa, and adult flies, making it possible to study pacing effects on a developing fly heart completely non-invasively.^3^

About 75% of human genes associated with disease have orthologues in *Drosophila.* In particular, *Drosophila* is a powerful genetic model system for cardiovascular studies, sharing many heart similarities with vertebrates at the early developmental stages. It has a short life cycle, and is cheap and easy to manipulate.^33, 34^ Additionally, *Drosophila* are not dependent on their heart tubes for oxygen transport and can survive for extended periods with severe defects. This capability makes *Drosophila* ideal candidates for longitudinal studies of congenital heart diseases which would be fatal at early life stages for other models. We have studied *Drosophila* cardiac function by noninvasively monitor the heart tube with optical coherence microscopy (OCM) in conjunction with heartrate modulation with optogenetic tools. OCM enables micron-scale 3D imaging of tissues in real time at imaging depths up to ∼400 um. The high resolution and high imaging speed allow for realtime monitoring of *Drosophila* heart function. The fly’s heart dimensions and heart rate can be directly measured from OCM videos to provide functional assessment of the *Drosophila* optogenetically controlled pacing.

In our previous work, we used blue light to pace *Drosophila* expressing ChR2. We were able to increase the heart rate of larva, early pupa and adult flies but were unable to reliably pace late pupa flies.^3^ The strong absorption of blue light within the darkened cuticle of late pupa flies prevented the excitation light from penetrating to the heart tube of late pupa. A few attempts have been made to address the light penetration problem, including surgical exposure of the heart tissue,^4, 35, 36^ optical fiber implantation,^4^ increasing the light intensity,^37^ and implanting μ-ILEDs.^38^ However, these methods are invasive and may damage the target and surrounding tissues. As an alternative to blue light, red light (> 600 nm) is a promising option for optogenetic stimulation of deep tissues.^37, 39^ Recently, a red-shifted excitatory opsin, ReaChR, was engineered by Lin et al. to activate neurons deep within the mouse brain.^40^ Unlike other opsins previously reported, this opsin exhibited high photocurrent efficiency at its broad spectral response peak centered on 600 nm. Inagaki et al. expressed ReaChR in the brain of *Drosophila* to control the complex behavior of a freely moving adult fly.^41^ Nyns et al. used a transgenic ReaChR rat model to terminate ventricular arrhythmias in an explanted whole heart, although they did not employ red-light stimulation.^35^ These studies suggest that incorporating ReaChR in the heart of *Drosophila* and performing red-light excitation can be a promising method to achieve non-invasive pacing through all the developmental stages. In this study, we seek to enhance the penetration depth of the stimulating light, extending *Drosophila* optogenetic pacing to all the developmental stages and improving the overall pacing efficiency.

Our previous experiment demonstrated excitatory pacing in which the *Drosophila* heart rate was increased. However, to model many heart conditions, it may be necessary to slow down or stop the *Drosophila* heartbeat. Halorhodopsin (NpHR), a widely used opsin with a red-shifted response spectrum, has been used to inhibit neuronal activities in different tissues through yellow-light stimulation.^2, 42–45^ By using red-shifted excitatory and inhibitory opsins, we can model a vast array of cardiac conditions, including tachycardia, bradycardia, and cardiac arrest. By inducing these cardiac conditions, we can study the cardiac fitness of these *Drosophila* models by assessing several metrics. For example, the cardiac recovery process after physical activity can be quantified by parameters such as the maximum heart rate during a physical challenge and the recovery time required to return to the resting heart rate after completing the challenge. In cardiac optogenetic pacing, the heart is forced to follow regulated frequencies, which could induce physiological cardiac distress. Investigating the heart rate recovery after optical stimulation could help to understand the impact of optical stimulation on heart function.

In this study, we demonstrated non-invasive red-light excitatory and inhibitory control of *Drosophila* cardiac function through cardiac optogenetics. Cardiac contractions were elicited or inhibited in two novel *Drosophila* organisms that had ReaChR or NpHR opsin transgenically expressed within the heart. The high penetration depth of red excitation light allowed us to successfully control the excitatory pace in late pupal flies. Red-light pacing had the added benefits of short excitation pulse widths and low stimulation power densities for successful cardiac pacing. We then demonstrated restorable cardiac arrest and inhibitory pacing in NpHR *Drosophila* for the first time. We also introduce several new metrics of cardiac fitness in *Drosophila*, such as the recovery period, or the maximum heart rate overshoot after induced cardiac arrest. The ability to perform noninvasive pacing and imaging can advance developmental studies of congenital cardiovascular defects, especially when combined with different transgenic fly systems.

## Results

Figure 1 provides a visual overview of this study. Figure 1a shows the custom red LED excitation and OCM imaging system used for simultaneous optogenetic pacing and OCM imaging. The red LED (∼ 617 nm) optically stimulates the fly heart. By using transgenic flies expressing ReaChR or NpHR in the heart, the light alters the conformation of the opsin and opens/closes the transmembrane region to allow ion flow and activate/inhibit heart contraction. The OCM non-invasively monitors the cardiac dynamics *in vivo* and in real time.^46^ Red light excitation requires lower amplitudes and shorter pulse widths than blue light to induce a heart contraction (Figure 1b).

**Figure 1.**
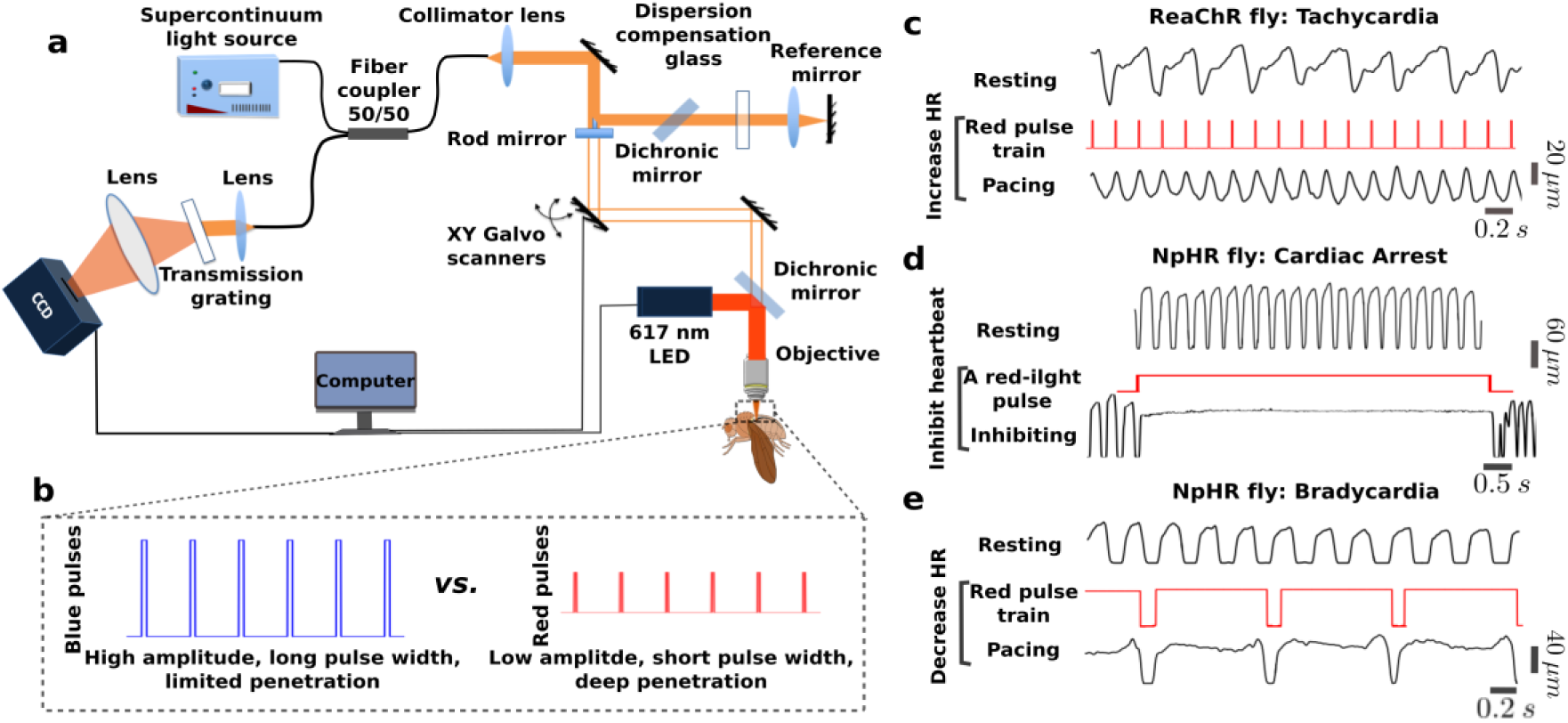
Red-light cardiac control in *Drosophila* expressing ReaChR or NpHR. **a,** Schematic illustration of the integrated red-light stimulation and OCM imaging system. **b,** Comparison of red-light and blue-light pulses, showing that red light has a lower amplitude, shorter pulse width, and deeper penetration depth for achieving cardiac pacing in *Drosophila melanogaster*. **c, d, e,** Three pacing strategies for red-light cardiac control. For each strategy, the upper trace shows the heart pulse change with time in the axial direction during rest. The middle trace shows the red-light pulse used for cardiac stimulation, and the lower trace shows heart pulse change with time in the axial direction with optical excitation. **c,** The heart pulse changes with time show an increasing heart rate (HR), mimicing tachycardia through red-light pacing of a ReaChR fly. **d,** Heart diameter changes with time to indicate inhibition of NpHR fly cardiac function for 10 s to mimic cardiac arrest through red-light excitation. **e,** Heart pulse changes with time to demonstrate decreasing HR of a NpHR fly to simulate bradycardia through red-light excitation.

We bred transgenic flies (UAS-ReaChR; 24B-GAL4 and UAS-NpHR; 24B-GAL4) in ten ReaChR and six NpHR fruit fly lines. The heart of *Drosophila* is close to the dorsal side and evolves with a tubular structure during development. Using our previously developed fly line screening protocol, described in the Methods section and in the Supplementary Materials, we found that ReaChR #53748 and NpHR #41752 were optimal fly stocks for excitatory and inhibitory pacing studies. Three pacing strategies were successfully demonstrated using the two fly stocks, as shown in Figure 1c-e. First, to simulate tachycardia, we increased the HR in transgenic ReaChR flies by pacing the heart with a red-light pulse train at a frequency higher than the resting heart rate (RHR) of the fly (Figure 1c). Second, to model cardiac arrest, we illuminated the heart of NpHR flies for 10 s locking open the cardiac ion channels (Figure 1d). After demonstrating cardiac arrest, we reduced the HR in NpHR flies to model bradycardia by illuminating the heart with red-light pulses at frequencies lower than the RHR (Figure 1e). In Figure 1c-e, for each simulation, we plotted the heart diameter over time in the resting and pacing states, with the excitation pulse shown in the middle.

### Optical simulation of tachycardia with ReaChR flies

We demonstrated red-light excitatory pacing of ReaChR *Drosophila* at the larval, early pupal, late pupal, and adult stages (Figure 2a-d, Supplementary video 1-4). As illustrated in the Figure, the heart was illuminated by red light pulse trains. Before optimizing the pacing pulse, a pulse width of 40 ms and excitation power density of 7.26 mW · mm^−2^ were used for pacing at each developmental stage. By analyzing 2D M-mode images acquired in 31 s through OCM imaging, we determined that the RHRs of the larval, early pupal, late pupal, and adult stages of the example fly were 4.4 Hz, 2.4 Hz, 2.5 Hz, and 5.7 Hz, respectively. Then, for example specimens of the same fly type as in Figure 2, we applied optical pacing at three frequencies above the initially measured RHR for all four stages: 5.5, 6.0, 6.5 Hz for the larval stage; 3.5, 4.0, 4.5 Hz for the early pupal stage; 4.5, 5, 5.5 Hz for the late pupal; and 9.0, 9.5, and 10 Hz for the adult stage. As shown in Figure 2a-d, the pacing rates (PRs) and measured heart rates (HRs) during pacing are indicated with red and black fonts in the M-mode images. The result demonstrated that each light pulse successfully elicited a heart contraction. The HRs followed the PRs, and reverted to the original individual RHRs after pacing was finished. The hearts of wild type (WT) control flies were not affected by the pacing light pulses at any developmental stage.

**Figure 2.**
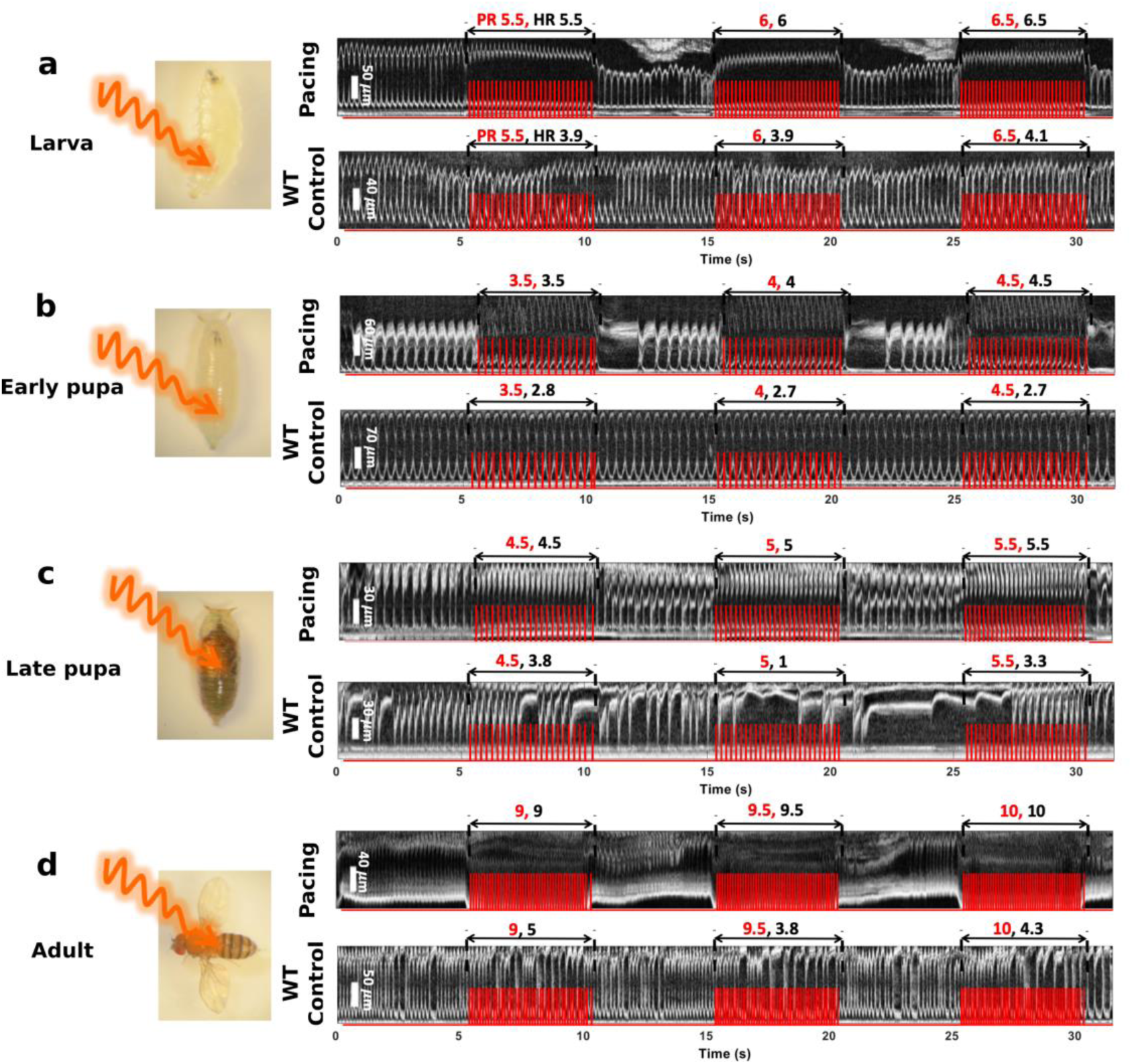
Optogenetic cardiac pacing of ReaChR-expressing *Drosophila* using red light at the larval, early pupal, late pupal, and adult developmental stages (see examples in Supplementary video 1-4). Demonstration of red-light optical stimulation of ReaChR-expressing *Drosophila* hearts, and 2D M-mode images acquired from ReaChR and WT flies with the OCM system during pacing at the larval (**a**), early pupal (**b**), late pupal (**c**), and adult (**d**) stages. The HRs of the ReaChR flies follow the PRs applied by the three different red-light pulse trains. M-mode images acquired from WT flies are shown as controls, in which applying the same pulse trains as for ReaChR flies had little effect on the fly heartbeat. Red traces in the M-mode images illustrate intensity changes in the excitation light. Red numbers in the M-mode images represent PRs, while black fonts indicate HRs during pacing.

After demonstrating red-light excitatory pacing, we optimized the pacing parameters, including the excitation power density and pulse width at the larval, early pupal, late pupal, and adult stages, as shown in Figure 3a-d. Figure 3a shows the pacing results for a late pupa, which are demonstrated for the first time. We varied the LED excitation power densities from 0.46 mW · mm^−2^ to 7.26 mW · mm^−2^ (top to bottom in Figure 3a) with a pulse width of 10 ms, and found that each pulse was able to elicit one heart contraction when the excitation power density was greater than 1.86 mW · mm^−2^. The HR diverged from the PR (5 Hz) at lower power densities. Statistical analysis of the power amplitude was performed by pacing multiple fruit flies at each stage (Figure 3b). The optimal red-light excitation power density of 3.63 mW · mm^−2^ enabled a hundred percent pacing probability at all the stages. In comparison, blue-light pacing required inputs of 35.5 mW · mm^−2^ for ChR2 larval and early pupal stages, and 12 mW · mm^−2^ for adults.^3^ The red-light optical exposure was well below the maximum permissible exposure (MPE) recommended by ANSI for safe use of lasers on skin (ANSI Z136.1) (7.26 mW · mm^−2^ *vs*. 27.41 mW · mm^−2^).

**Figure 3.**
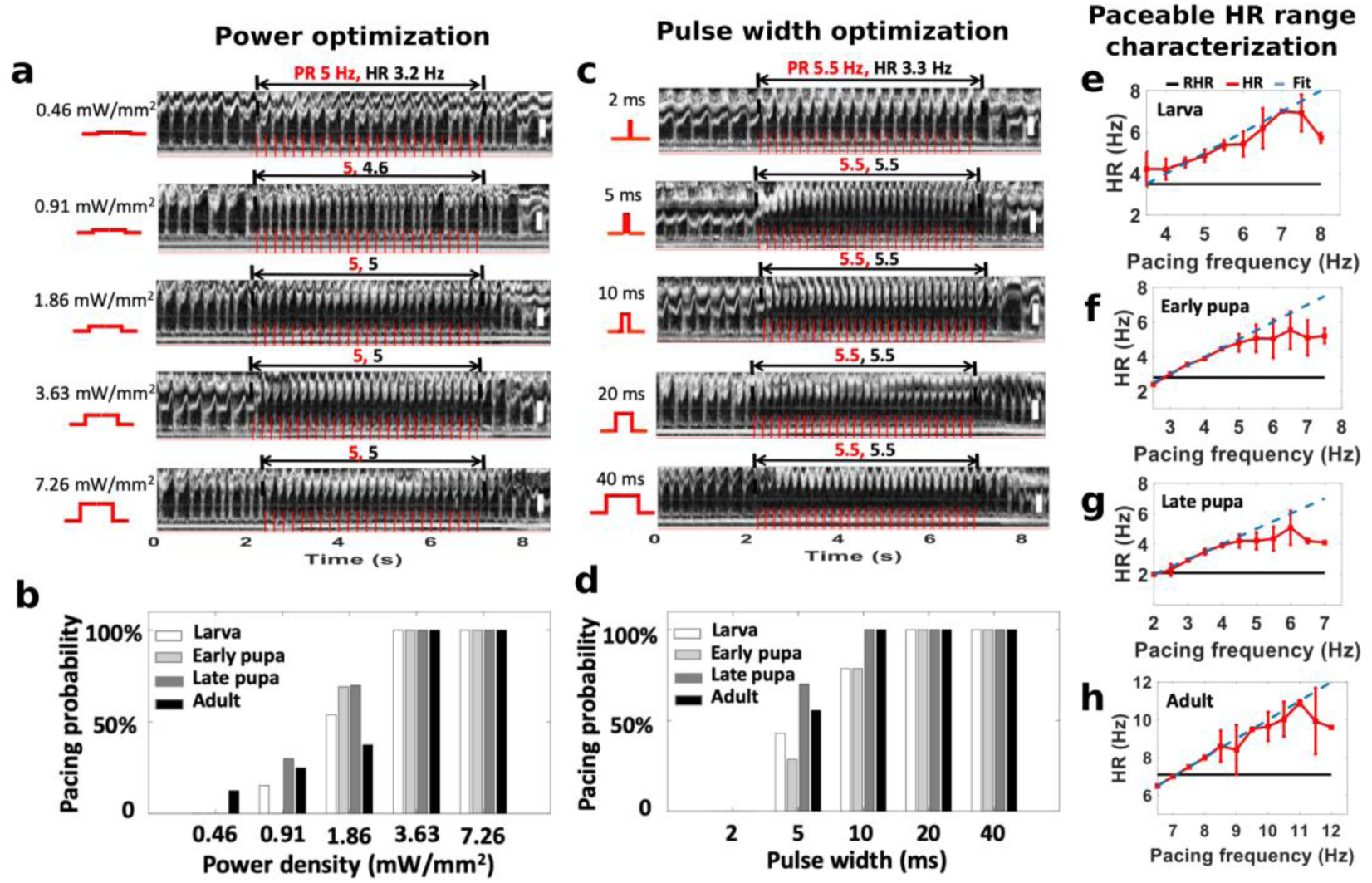
Finding the optimal excitation power density and pulse width for red-light optogenetic pacing of ReaChR-expressing *Drosophila*. **a,** M-mode images acquired from a late pupa with a resting heart rate (RHR) ∼3 Hz, showing the HR following a pacing rate (PR) of 5 Hz with pulse intensities tuned from 1.86 to 7.26 mW · mm^−2^, and the HR deviation from the PR with pulse densities lower than 0.91 mW · mm^−2^. **b,** The effect of excitation power densities on pacing probabilities for larva (n = 14), early pupae (n = 17), late pupae (n = 11), and adult (n = 12) stages, showing that decreased excitation power densities lower than 3.63 mW · mm^−2^ led to lower pacing probabilities. **c,** 2D M-mode images acquired from a late pupa with an RHR ∼2.5 Hz, showing the heartbeat following the pacing rate (PR) of 5.5 Hz as the pulse widths are tuned from 5 ms to 40 ms, and the heart rate (HR) deviation from the PR with a pulse width of 2 ms. **d,** The influence of stimulation pulse width on pacing probabilities for larva (n = 14), early pupa (n = 17), late pupa (n = 11), and adult (n = 12) stages. Pulse widths of 10 ms or less resulted in lower pacing probabilities for the first two stages. Width of 5 ms lowered the pacing probabilities for the late two stages. **e-h,** Plots of the average RHR (black solid line), the average HRs from tuning the pacing frequencies (red curve), and the fitting line of the average HRs (blue dashed line) characterizing the HR adjustable ranges for larva (n = 14), early pupa (n = 16), late pupa (n = 11), and adult (n = 10). The HR was tuned around the average RHR of the respective stages. In larvae, the fly heart linearly followed the pacing frequency from 3.5 to 5.5 Hz, where the average RHR was 3.5 Hz. In early pupae, the average RHR was 2.8 Hz, and the heart linearly followed the pacing frequency in the range between 2 and 5 Hz. For late pupae, the average RHR was 2.1 Hz, and the heart linearly followed the pacing frequency in the range between 2.4 and 4.5 Hz. During the adult stage (with a higher average RHR of 7 Hz), the HR and pacing frequency showed a linear relationship between 7 and 8.5 Hz. The fly heart reliably followed the pacing frequency up to ∼167%, ∼218%, ∼219% and ∼143% of the RHR at the larva, early pupa, late pupa and adult stages, respectively. Results are presented as mean ± SD. Red traces on the M-mode images illustrate the intensity change of excitation light. Red and black fonts in M-mode images represent PRs and HRs respectively, as in Figure 2. Scale bars throughout the figure, 30 μm.

With the optimized power density, we further characterized the optimal excitation pulse width. The M-mode images in Figure 3c illustrate the HRs of a late pupa paced with pulse widths of 2 ms, 5 ms, 10 ms, 20 ms, and 40 ms (top to bottom). Pulse widths wider than 5 ms led to robust optical pacing. The influence of the pulse width on the pacing probability was then quantified for each developmental stage (larva, early pupa, late pupa, and adult) (Figure 3d). At each stage, a hundred percent pacing probability could be achieved with a pulse width of 40 ms. The minimum pulse widths for this achievement were demonstrated to be 20 ms for the larval and early pupal stages, and 10 ms for the late pupal and adult stages. As expected, shorter pulse widths resulted in lower success rates at each stage. No flies could be paced when the pulse width was decreased to 2 ms.

With the optimal power density and pulse width, the paceable frequency range over which pacing could be achieved was studied for individual developmental stages (Figure 3e, f, g, h). The maximum paceable heart rates (maxPR) were observed to be ∼167%, ∼218%, ∼219%, and ∼143% of the RHR for the larval to adult stages, demonstrating a broad controllable HR range throughout the life cycle of *Drosophila*. Pacing frequencies too far from the RHR caused the HR to deviate from the PR, which is consistent with previous conclusions obtained using ChR2.^2, 3^

### Optically modelling cardiac arrest and bradycardia

#### Restorable cardiac arrest

With NpHR flies, we demonstrated restorable cardiac arrest at different developmental stages (Supplementary video 5-7). The inhibitory microbial opsin NpHR was expressed in the heart of *Drosophila* along with YFP through the transgenesis technique (Figure 4a). As shown in Figure 4b, the M-mode images indicate the influence of continuous red-light illumination on the heart functions. The heartbeats of the larval, early pupal, and late pupal flies instantly stopped when illuminated with red light. Cardiac arrest was sustained during continuous illumination of the heart for 10 s. After switching off the red light, the heartbeat immediately restarted. A control experiment was performed using WT flies for each stage. The heart function was unaffected by red light stimulation in control flies.

**Figure 4.**
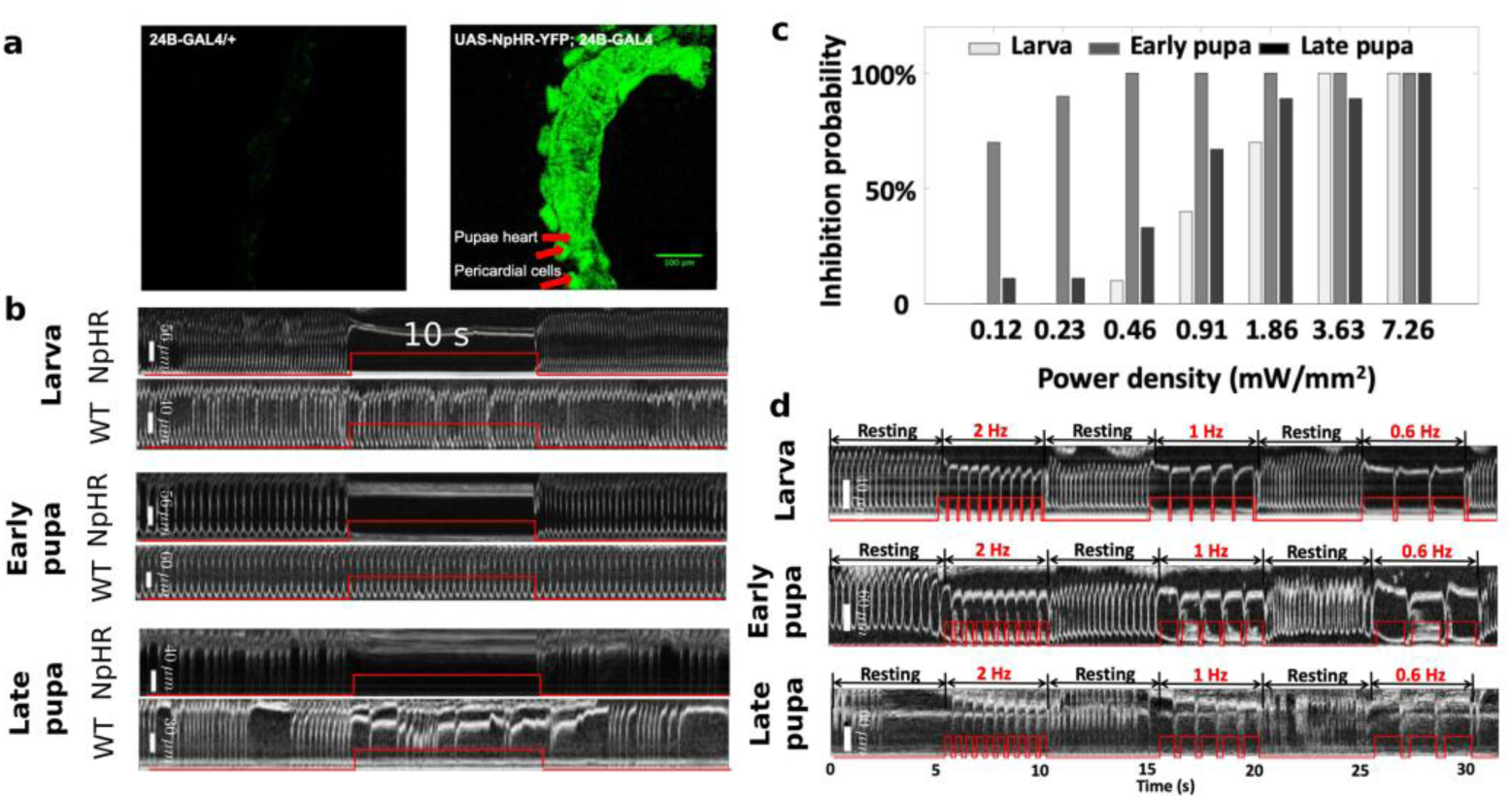
Inhibition of cardiac function to mimic cardiac arrest and bradycardia. **a,** Fluorescence images showing strong expression of YFP (co-expressed with NpHR) in the heart of a pupal fly. **b,** The hearts of NpHR and WT flies were illuminated constantly for 10 s using red light at the larval, early pupal, and late pupal stages, respectively (see examples in Supplementary video 5-7). The heartbeats of the NpHR flies stopped immediately when red light illumination was applied. The cardiac arrests lasted for 10 s, and regular heartbeats restarted after the light was turned off. M-mode images acquired from WT flies at the three developmental stages are shown as control. Little cardiac function alteration was observed by applying the same constant red-light illumination as for NpHR flies. **c,** Characterization of the inhibition probabilities for different power densities of the stimulation light. **d,** M-mode images showing inhibitory pacing of NpHR flies with three red-light pulse trains at frequencies of 2 Hz, 1 Hz, and 0.6 Hz respectively at larval, early pupal, and late pupal stages (see examples in Supplementary video 8-10). To achieve successful inhibitory pacing, the pulse width was tuned with the given excitation frequency for each pulse train. The heart rates of the fruit flies at the three stages were successfully reduced to 2 Hz, 1 Hz, and 0.6 Hz, even though the RHRs are different. Similar to the cardiac arrest simulation, the heart resumed beating regularly after optical pacing was terminated. Red traces on the M-mode images illustrate the intensity change of excitation light. Red and black fonts in M-mode images represent PRs and HRs respectively, as in Figure 2.

We characterized the red-light excitation power density to study the probability of inducing cardiac arrest at different stages (Figure 4c). The power density was tuned up from 0.12 mW · mm^−2^ to 7.26 mW · mm^−2^, and applied to larval, early pupal, and late pupal flies. Multiple flies were measured for each stage as seen in Supplementary Table 4. For each stage, 3.63 mW · mm^−2^, 0.46 mW · mm^−2^, and 7.26 mW · mm^−2^ were the lowest powers capable of efficient cardiac inhibition. Lower power levels led to reduced cardiac arrest probabilities. In particular, early pupal flies required relatively low power (*e.g.* 0.46 mW · mm^−2^) for reliable cardiac arrest.

#### Restorable bradycardia

With the ability to suppress cardiac contractions in NpHR flies, it is possible to simulate the regularly seen cardiac arrhythmia, bradycardia (Supplementary video 8-10). Here, we optimized optogenetic inhibitory pacing by simultaneously tuning the frequency of the excitation-light pulse train and the pulse duty cycle for each NpHR fly.

For a specific frequency, the duty cycle was tuned to allow one heart contraction between two adjacent pulses. As shown in Figure 4d, we controlled the HRs of the example NpHR larva, early pupa, and late pupa by adjusting the duty cycle for individual red-light pulse trains at frequencies of 2 Hz, 1 Hz, and 0.6 Hz, all at a power density 7.12 mW · mm^−2^. For the larva, duty cycles of 80%, 90%, and 90% were used for the respective frequencies. The time between adjacent pulses in each pulse train, 100 ms, 100 ms, and 167 ms, allowed for one heart contraction by the larva at each frequency. For the early pupa, inter-pulse times of 175 ms, 200 ms, and 333 ms were used to allow for a single heartbeat, and 200 ms, 300 ms, and 333 ms were used for the late pupa. The heart rates were successfully slowed to various lower frequencies, and restored to the RHR after the optical stimulation was suspended. Even with different RHRs, the larva, early pupae, and late pupae could be paced at identical frequencies.

### Heart rate recovery after cardiac arrest

During simulation of cardiac arrest using NpHR flies, we observed heart rate recovery process after red-light stimulating the fly heart. We investigated the recovery dynamics using NpHR early pupal flies by inducing cardiac arrest for different time periods. Specifically, the heart of an NpHR early pupa was monitored for 60 s, with cardiac arrest induced after imaging the heart for 5 s in the resting state. To induce different cardiac arrest periods, red-light LED illumination was continuously applied for periods of 1 s, 2 s, 5 s, 10 s, and 20 s respectively for each fly. Heart rate recovery after the illumination was studied by analyzing the maximum heart rate (maxHR) and the recovery time. The heart rate was normalized by the mean RHR of the first 5 s of the OCM recording to adjust for the heart rate differences between individual flies. We regard a normalized heart rate within 80%-120% as a restored RHR after cardiac arrest. Figure 5a shows the dynamic heart rate recovery process of an NpHR early pupa after cardiac arrest challenges of 1 s, 2 s, 5 s, 10 s, and 20 s, respectively. With 1 s and 2 s cardiac arrest, the fly heart rate returned to its RHR immediately after red-light inhibition. With the longer inhibition time, an overshoot of HR and an increased recovery time were observed following the cardiac arrests. With a 20 s inhibition, an ∼150% HR overshoot and ∼25.4 s recovery time were observed from this fly. To further evaluate the consistency of this phenomenon, we induced cardiac arrest in 18 flies for different stimulation durations and imaged the fly heart beat for 60 s to analyze the heart rate recovery. In Figure 5b, we plotted the mean and standard deviation of the heart rate at each time point. Consistent with what is shown in Figure 5a, the fly HR returned to the RHR immediately after 1 or 2 s of cardiac arrest stimulation. For longer stimulations, the mean HR overshoot and recovery time were found to be 130% and 4.62 s, 141% and 8.17 s, and 177% and 10.85 s for cardiac arrest times of 5s, 10s, and 20 s, respectively. To look into the relationship of the cardiac recovery process with arrest time, we compared the means and medians of the HR overshoots and recovery times for the five cardiac arrest periods. As shown in Fig. 5c, d, the overshoot and recovery time started to show significant differences when the arrest time increased to 5 s.

**Figure 5.**
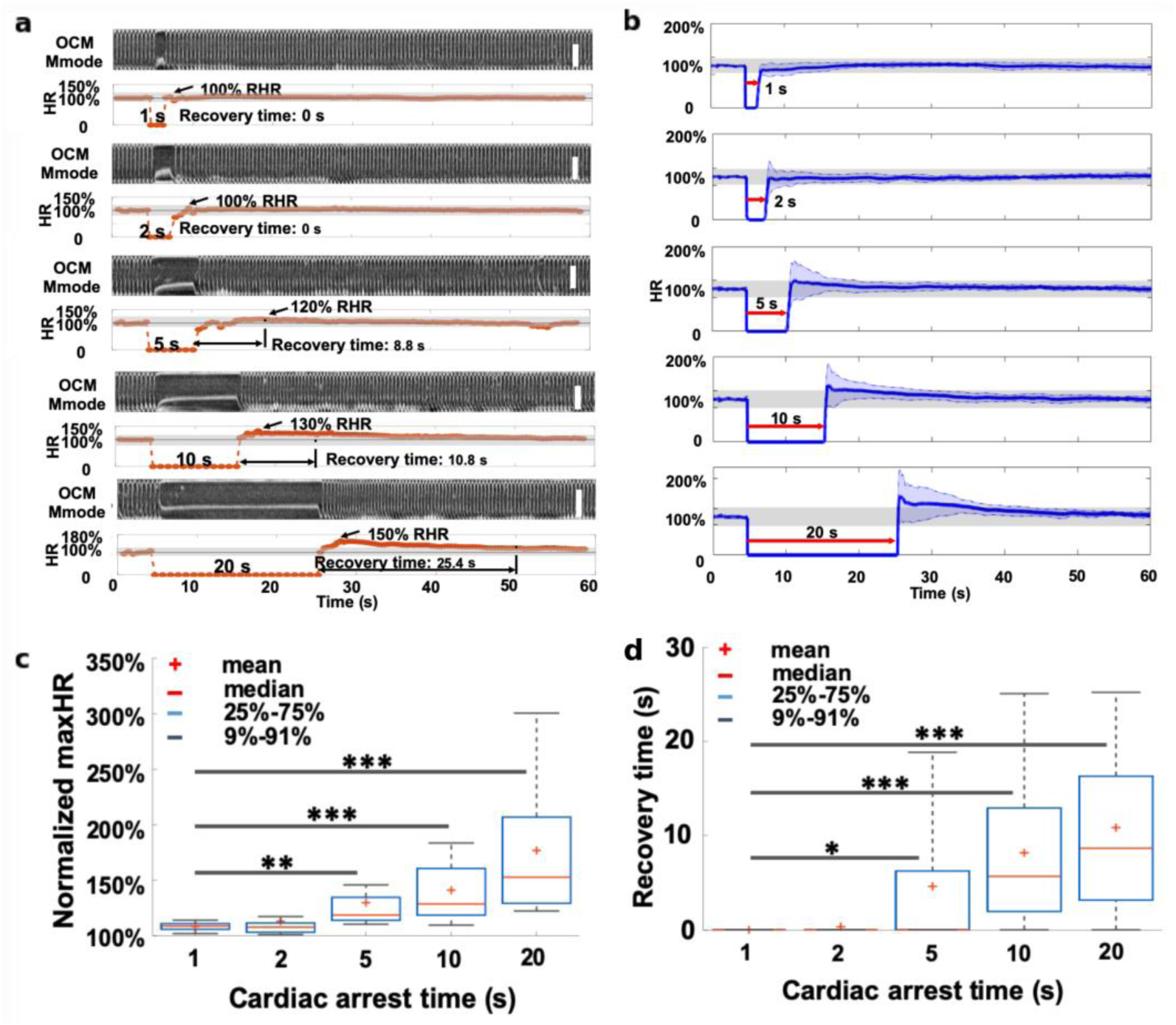
Heart rate recovery after pacing induced cardiac arrest in NpHR early pupa. **a,** Demonstration of heart rate recovery after red-light LED excitation for 1 s, 2 s, 5 s, 10 s, and 20 s, respectively in an NpHR early pupa (see examples in Supplementary video 11-15). An increased overshoot and recovery time were observed with longer cardiac arrest times. Scale bar: 120 μm. **b,** Group average of heart rate change after creating cardiac arrest for 1 s, 2 s, 5 s, 10 s, and 20 s, using red-light excitation (n = 18). The blue shade represents the standard deviation of the heart rate at each time point. The gray bar represents the normalized heart rate within 80% - 120% of the RHR. Characterization of the mean, median, 25^th^ percentile, and extremes for different cardiac-arrest-induced normalized maximum heart rates (**c**) and recovery times (**d**). Two-sided Student’s t-tests were used. *, p < 0.05; **, p < 0.01; ***, p < 0.001.

## Discussion

Light in the red to infrared range undergoes low scattering and absorption.^41^ Multiple red-shifted opsins have been engineered in the past several years to excite or inhibit neural or cardiomyocyte activities.^40, 47–50^ Some red-shifted opsins have been applied in optogenetic control of deep tissue, such as controlling neuronal activities in the mouse cortex through an intact skull,^40^ and behavioral control of freely moving flies.^41, 51^ Here, we leverage the features of red-shifted opsins to demonstrate non-invasive cardiac control through red-light stimulation.

Among the available red-shifted excitatory opsins, ReaChR is popular with optogeneticists for applications requiring deep penetration.^35, 41, 51^ In this study, we applied ReaChR to the cardiac *Drosophila* model. We successfully showed that red-light stimulation can induce depolarization in ReaChR-expressing *Drosophila* hearts, causing heart contractions. Previously, late pupae expressing ChR2 were not able to be paced due to their highly pigmented cuticle,^3^ but red light has been confirmed to have deeper optical penetration through the cuticle of *Drosophila* than blue light.^41^ Here, in addition to pacing larva, early pupae, and adult flies, the late pupae were also able to be paced successfully. With the addition of late pupae, optogenetic tachypacing can be successfully carried out throughout the *Drosophila* life cycle.

Tachypacing ReaChR-expressing flies was further explored by looking into the excitation power density, pulse width, and extensible heart rate. Pacing probabilities decreased with reduced excitation power densities, which was expected since lower power leads to decreased photocurrent passing through myocardial cell membranes and is less likely to open ReaChR-constructed ion channels. The minimum power density for robust pacing with red light was demonstrated to be 3.63 mW · mm^−2^, which is considerably lower than the required blue light energies of 35.5 mW · mm^−2^ for larvae and early pupae, and 12 mW · mm^−2^ for adults.^3^ The increased efficiency may be accounted for by the enhanced penetration depth of red light. ReaChR flies also required shorter illumination pulse widths for successful excitation than ChR2-expressing flies. Here, 20 ms was demonstrated as the optimal pulse width for pacing larval and early pupal flies, while only 10 ms was required for pacing late pupal and adult ReaChR flies. Adult ReaChR flies were paced more efficiently than ChR2 adults, which required 20 ms for reliable pacing. Furthermore, 100% of ReaChR adult flies were able to be paced, indicating high pacing reliability.^3^ The short optimal pulse widths required for excitatory pacing may be caused by the fast kinetics of ReaChR-constructed cation channels.^3, 40^ Fast opsin kinetics indicate rapid opening of the light-gated channels under illumination, which allows sufficient cations to cross the cell membrane and elicit action potentials in cardiomyocytes.^40, 52^

To model pathologies such as ventricular arrhythmia or bradycardia, we developed *Drosophila* models expressing the inhibitory opsin NpHR within the myocardium.^35, 36^ Here, we are able to use NpHR-expressing *Drosophila* to reliably mimic both cardiac arrest and bradypacing in the larval, early pupal, and late pupal stages. This is the first demonstration of restorable cardiac arrest and precise control of heartrates less than the RHR in intact *Drosophila* at multiple developmental stages. In the adult stage, cardiac function inhibition was observed only occasionally and for only 1-2 s in some flies (Supplementary Figure 1). This phenomenon might be due to a more active metabolism or a more extensively innervated adult cardiac muscle layer. High inhibition probabilities were demonstrated at low excitation power densities in the larval, early pupal, and late pupal stages. The optimal power densities were demonstrated to be 3.63 mW · mm^−2^, 0.46 mW · mm^−2^, and 7.26 mW · mm^−2^ respectively for each stage, all of which are less than the ANSI standard for skin exposure, indicating safe LED red-light stimulations.

Our new research platform not only enables simulation of various cardiac pathologies, but also enables investigation of cardiac physiology after a short challenge. For humans, the heart rate and recovery time after exercise are good indicators of fitness. Health problems lead to higher heart rates and prolonged recovery periods after exercise. In previously exercised *Drosophila*, the recovery from cardiac arrest induced by electrical pacing was shown to be faster than the recovery period for *Drosophila* that had not been exercised^53^. Here the arrest recovery dynamics were initiated through optogenetic stimulation of NpHR *Drosophila*. The heart rate rapidly overshot the RHR after optical stimulation, and then exhibited exponential decay until returning to the resting heartrate. This recovery paradigm is similar to heartrate recovery after physical exercise in humans. The maximum heart rate and recovery time were shown to increase with arrest duration, suggesting optogenetic-stimulation induced indicators of cardiac health common to both vertebrate and non-vertebrate models.

While cardiac pacing’s most famous and beneficial application, implanted electrical pacemakers in humans, is clinical, cardiac pacing can also be used as a powerful research tool. Electrical pacing in *Drosophila* has been used extensively to quantify aging-induced cardiac dysfunction and arrhythmias.^54–56^ Studies have induced cardiac decline in *Drosophila* with electrical tachypacing and have correlated the organisms’ susceptibility to this decline with aging, and as well as associating it with human ortholog disease genes.^57–59^ To tachypace the *Drosophila* in these studies, electrical current was applied across the entire fly, which causes inhomogeneous and non-specific stimulation and may cause thermal damage or other detrimental effects to the animal. Here, we not only demonstrate tachycardia effectively and non-invasively in *Drosophila*, but also are able to conduct real-time imaging for heart function monitoring. With the non-invasive platform, each adult can be longitudinally tracked during the aging process, without requiring that each fly be sacrificed after each pacing performance, which is promising for aging-related heart function studies.

More extensive research could be performed based on this study. First of all, new opportunities could be explored in cardiac pathology studies. In vertebrate models, including rats, rabbits, and dogs, cardiac pacing has been used to simulate tachycardia, bradycardia, and cardiac failure, and to study molecular and epigenetic effects of these cardiac modes. With tachycardia animal models, dynamics such as conduction velocity and gap junction properties,^60^ hemodynamic deterioration,^61^ and intracellular Ca2+ dynamics^62^ have been analyzed. Potassium channel subunit regulation,^63^ and the relation between endogenous physiological late Na+ current and bradycardia-related ventricular arrhythmias^64^ were investigated by simulating bradycardia. ReaChR and NpHR could be expressed in these vertebrate models to study these long term epigenetic changes with reduced invasiveness over electrical pacing, although, with vertebrates, surgery would still be required to implant illumination devices. Second, the developed red-shifted fly models could be complementary to vertebrate models for physiological or pathological studies via red light cardiac control. They could also be further transgenically manipulated to study human-ortholog-gene mutant induced pathologies.

The OCM integrated platform also provides new opportunities in both hardware and software for animal cardiac studies. The non-invasive nature of the OCM technique enables longitudinal monitoring of cardiac function during optical control without damaging the cardiac tissue. As a label-free technique, it could become a promising complement to previous *Drosophila* cardiac assessment methods that require staining, such as optical mapping, immunofluorescence, or histology.^5, 36^ The use of red excitation light for optogenetic cardiac control enables higher penetration depth and longitudinal cardiac arrhythmia studies. The combination of OCM and optogenetics establishes an all-optical, non-invasive manipulation and real-time imaging modality for cardiac optogenetics.

Even though our system is promising for cardiac studies, there are several limitations. First, the LED light source used for red-light excitation is difficult to focus specifically on the pacemaker region in *Drosophila*, which increases the total power necessary for excitation. This concern could be addressed by using a red laser source instead of an LED. Second, even though efficient non-invasive cardiac control was realized in *Drosophila* using red LED light, the penetration depth is insufficient for controlling the heart functions of intact large animals. The OCM system is also not suitable for large animal heart imaging *in vivo*.

## Conclusions

In this study, we developed ReaChR and NpHR *Drosophila* models and an integrated non-invasive red-light cardiac control and OCM imaging system. We successfully mimicked restorable tachycardia, bradycardia, and cardiac arrest at different developmental stages, and we demonstrated improved pacing efficiencies using red LED light (617 nm) stimulation. The *Drosophila* heartbeat can be precisely manipulated in real time by modifying the frequency and pulse width of the red light. The recovery dynamics after red light stimulation of NpHR flies were observed and quantified. The red-light stimulation, OCM imaging, and transgenic *Drosophila* systems provide promising research platforms for cardiac optogenetic studies in the future.

## Methods

### Transgenic fly model and fly culture

24B-GAL4 was used as the driver of UAS-mediated opsin expression^3^ in the heart of *Drosophila melanogaster*. By crossing UAS-ReaChR or UAS-NpHR-YFP flies with 24B-GAL4 flies, targeted expressions of ReaChR (UAS-ReaChR; 24B-GAL4) and NpHR (UAS-NpHR; 24B-GAL4) were achieved in cardiac tissue. Wild type (WT) W118 flies were crossed with 24B-GAL4 flies to obtain 24B-GAL4/+ as the control.

We prepared fly food by mixing Formula 4-24 (Instant *Drosophila* Medium; Carolina Biological Supply Company), water, and 10 mM all-*trans*-retinal (ATR) (Toronto Research Chemicals Inc.) solution, which used 200 proof ethanol as the solvent. The ATR concentrations in food used for culturing ReaChR and NpHR flies were 3 mM and 10 mM, respectively, in order to achieve sufficient opsin expressions in the heart. Developed flies were transferred into a vial with the prepared formula and kept in an incubator at 25 °C for ∼10 h for cross breeding. Flies in the first generation were obtained at the larval, early pupal, late pupal, and adult stages for cardiac control and optical imaging experiments. Normal food was prepared by mixing formula 4-24 and water for culturing WT flies at the same temperature.

### Integrated red-light excitation and OCM imaging system

We developed an integrated red LED excitation and OCM imaging system for simultaneous, non-invasive cardiac stimulation and monitoring in *Drosophila* (Figure 1a). The OCM system used a portion of a supercontinuum laser source to provide near-infrared light with a central wavelength of 800 nm and a bandwidth of 220 nm.^3^ The axial and transverse resolutions of ∼1.5 μm and ∼3.6 μm in tissue were obtained by using the broadband source and a 10x objective lens respectively to resolve the fly heart structure. A 2048 pixel line scan camera was utilized to detect the interference signal of the back-reflected light from the two arms of the OCM system.

To perform optogenetic control of heart functioning, we integrated a 617 nm LED light source (Migtex Systems, BLS-LCS-0617-03-22) into the OCM sample arm through a dichroic mirror and focused the light to the same point of the imaging beam through the objective lens with a spot size of ∼2.2 mm, covering the entire heart tube. Control software provided by Migtex Systems allowed tuning of the light intensity. A Labview program was used to control the pulse width (or duty cycle) and frequency of the LED signal, and to synchronize the M-mode image acquisition for the OCM system through a function generator (Agilent 33210A, Keysight technologies, USA) and a DAQ card (National Instrument USB-6008).

### Optical excitation and OCM imaging protocol

The focused excitation light illuminated the entire heart tube in both ReaChR and NpHR flies at each developmental stage. For excitatory optical pacing, power densities of 7.26 mW · mm^−2^ and 3.63 mW · mm^−2^, and pulse widths of 20 ms and 10 ms were used for larva and early pupa screenings, and late pupa and adult fly screenings, respectively, to identify the flies which could be paced. Flies which could be successfully paced at frequencies 20% higher than their RHRs were selected for experiments. Three pacing pulse trains were applied serially during each OCM measurement, with each pulse train lasting for ∼4.5 s. Then, 4096 cross-sectional images were acquired in ∼32 s at the frame rate of ∼130 frames/s from the posterior segments of the larval and early pupal heart, and the anterior portions of the late pupal and adult heart. M-mode images were acquired from the cross-sectional images, using ImageJ to analyze the time-lapsed heart function with and without red-light stimulations. Control flies were paced and imaged with identical procedures. For all the identified ReaChR flies, we modulated the excitation power densities from 0.46 mW · mm^−2^ to 7.26 mW · mm^−2^, the pulse widths from 2 ms to 40 ms, and pulse frequencies from 0.5 Hz above the RHRs to higher values until divergence of HR from PR was observed.

### Cardiac arrest and inhibitory pacing protocol

To demonstrate restorable cardiac arrest, 10 s of continuous red light was used to identify viable larva, early pupa, late pupa, and adult flies, with an excitation power density of 7.26 mW · mm^−2^. After discerning viable flies, power intensities were tuned from 0.12 mW · mm^−2^ to 7.26 mW · mm^−2^ to characterize the effect of stimulation power on the cardiac arrest rate for each developmental stage. For inhibitory pacing, duty cycles of excitation pulse trains were modified for individual flies. Pulse frequencies of 2 Hz, 1 Hz, and 0.6 Hz were tested at the excitation power density of 7.26 mW · mm^−2^. The single heart contractions between adjacent pulses indicated the heart rate at the respective frequencies.

### Protocol for study of heart rate recovery

To investigate the heartbeat recovery after cardiac arrest in NpHR flies, we monitored the heart of each early pupa for 60 s through the OCM imaging technique. Before demonstrating cardiac arrest, the minimum excitation power was determined for individual flies by observing the heartbeat during red light excitation. After allowing the heart to beat for 5 s at the RHR, different cardiac arrests were then generated by stimulating the heart for 1 s, 2 s, 5 s, 10 s, or 20 s. To compare the cardiac-arrest-induced cardiac recovery dynamics between different arrest periods, two-sided Student’s t-tests were used for statistical analysis and results were considered statistically significant at p < 0.05.

### Data availability

The data and code that support the findings of this study are available from the corresponding author upon reasonable request.

## Acknowledgements

We would like to acknowledge our funding support by the National Science Foundation [NSF1455613 to C.Z. and A.L., the National Institute of Health (R15EB019704 to C.Z. and A.L., R01EB025209 to C.Z., R03AR063271 to A. L and R01AG014713 and R01MH60009 to R.E.T.), and Cure Alzheimer’s Fund [to R.E.T.]. We would also like to acknowledge J. Ballard, G. Ni, Q. Guo, S. Wang, J. Yu, F. Cai, Z. Li and Y. Huang for suggestions of figure preparation and helpful discussions.

## Author contributions

C.Z. and J.M. conceived and designed the experiments; A.L. and R.E.T. provided access to the fly facilities; A.L. and Z.L. prepared the fly models; J.M. performed the experiments and analyzed the data; J.M., J.J., C.Z., and A.L. wrote the manuscript. All authors read the manuscript and provided constructive feedbacks.

## Supplementary materials

### I. Fly Line Development and Assessment

#### A. Characterization of fruit fly lines for red-light excitatory pacing

We developed fly models for red light excitatory pacing by crossing 24B-GAL4 with UAS-ReaChR and UAS-CsChrimson flies obtained from Bloomington Drosophila Stock Center, including fly lines #53740, #53741, #53742, #53746, #53747, #53748, #53749, #55134, #55135, and #55136 (Supplementary Table 1). The red-shifted microbial opsins ReaChR and CsChrimson were targeted for cardiac tissue expression through UAS mediation and the 24B-GAL4 driver system. ReaChR and CsChrimson flies were mated and cultured on Formula 4-24 (Instant Drosophila Medium; Carolina Biological Supply Company) with all-*trans*-retinal (ATR) (Toronto Research Chemicals Inc.). The first generation of each fly stock was cultured at 25 ℃ and tested by red light optogenetic pacing in the larval, early pupal, and adult stages. M-mode images obtained with the OCM system showed no change in heart function for the #53740, #53741, #53742, #53746, and #53747 flies, indicating unsuccessful pacing. CsChrimson #55134 flies could be paced in the larval and pupal stages, but only female flies could be successfully paced. Pacing was successfully performed in the larval and pupal stages of the #53749, #55135, and #55136 stocks, but the flies were not able to undergo eclosion. A heartbeat pause was also observed in #53749 larval flies after red light stimulation. The heart restarted beating after ∼5 minutes. The first generation of the ReaChR #53748 fly stock were able to develop into adults and were able to be paced throughout their lifecycle. This optimized fly stock was used to demonstrate excitatory pacing and simulate tachycardia.

**Supplementary Table 1.**
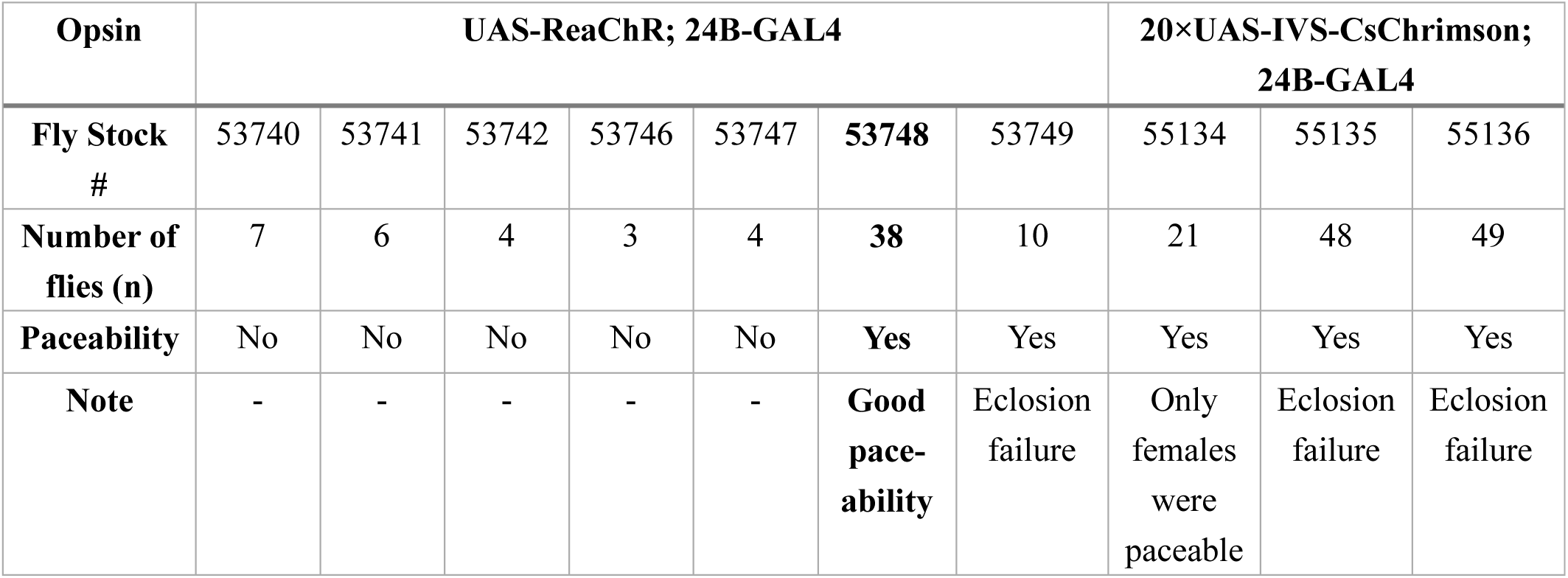
Characterization of fruit fly lines for red-light optogenetic pacing, listing the fly stock and number of screened flies for UAS-ReaChR; 24B-GAL4 and 20×UAS-IVS-CsChrimson; 24B-GAL4 flies respectively. All flies were cultured at 25 °C. No: unable to be paced; Yes: able to be paced.

#### B. Characterization of fruit fly lines for simulating cardiac arrest

The UAS-NpHR-YFP; 24B-GAL4 transgenic fruit fly models were engineered by crossing the UAS-NpHR-YFP fly from Bloomington Drosophila Stock Center (#41752) with 24B-GAL4 flies. The same culturing protocol as previously described was used, but with an ATR concentration of 10 mM to ensure sufficient expression of NpHR in cardiomyocytes. The first generation flies were evaluated at the larval stage by stimulating the heart for 4 s at a power density of 7.26 mW/mm^2^. Heart beat inactivation was observed in ten flies in response to the red-light illumination. The UAS-NpHR-YFP; 24B-GAL4 flies were then used for mimicking cardiac arrest and bradycardia.

Constant red-light stimulation was applied to the larva, early pupa, late pupa, and adult NpHR flies. Inhibition of cardiac function was observed in the first three developmental stages. In UAS-NpHR-YFP; 24B-GAL4 adult flies, red light stimulation could cause the heart to pause for 1 to 2 seconds (Supplementary Figure 1a), but WT adults had no response to the red-light stimulation (Supplementary Figure 1b). The fly heart undergoes significant remodeling during metamorphosis, which might affect the NpHR expression level, leading to partial failure of cardiac inhibition in adult flies. The high metabolic level of adult fly heart may also prohibit it from stopping for over a few seconds with red light inhibition.

**Supplementary Figure 1.**
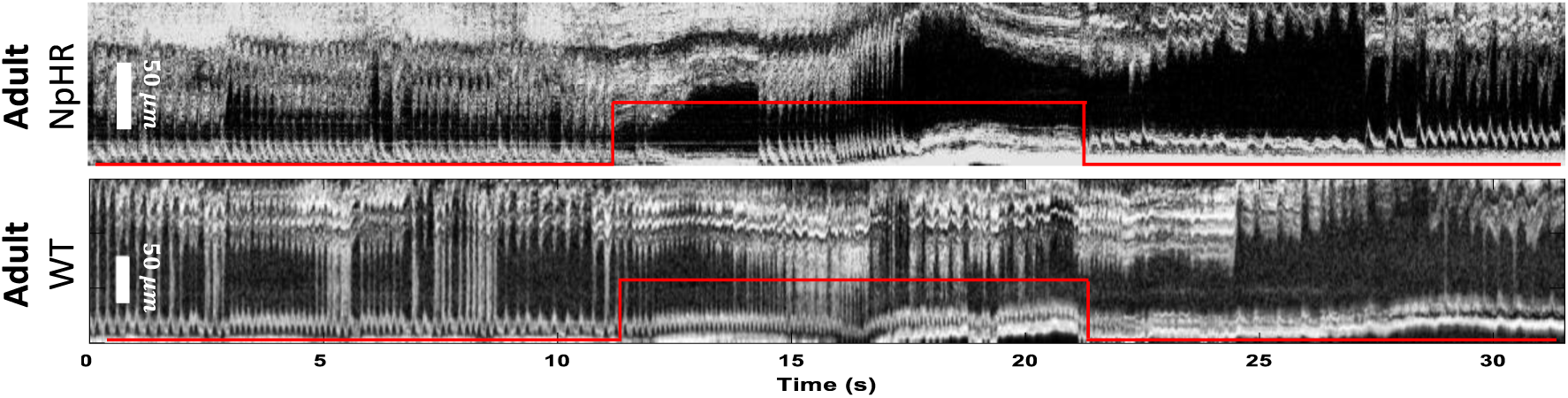
M-mode images demonstrating inhibition of cardiac function with 10 s constant red-light stimulation of NpHR and WT adult flies. Short term inhibition of cardiac function was observed in NpHR flies after the red light was turned on, while no response to red light stimulation was observed within WT flies.

### II. ReaChR flies used for characterization of red-light pulses for excitatory pacing

To characterize excitation pulses for successfully pacing ReaChR flies, the excitation power density, pulse width, and pacing frequency were investigated at each developmental stage. ReaChR larva, early pupa, late pupa, and adult flies were paced and imaged using our integrated red LED and OCM system. The flies studied are listed in Supplementary Table 2.

**Supplementary Table 2.** The number of UAS-ReaChR; 24B-GAL4 flies measured at different developmental stages for characterization of pacing pulse width, power density, and pacing frequencies.

| Parameters/developmental stages | Larva | Early pupa | Late pupa | Adult |
| --- | --- | --- | --- | --- |
| Optimize pulse width (n) | 14 | 17 | 11 | 12 |
| Optimize power density (n) | 14 | 17 | 11 | 12 |
| Characterize pacing frequency (n) | 14 | 16 | 11 | 10 |

### III. NpHR flies used for characterization of the excitation power density needed to mimic cardiac arrest

The correlation of the cardiac arrest probability with excitation power density was determined by tuning the red-light power from 0.12 mW/mm^2^ to 7.26 mW/mm^2^. A cardiac arrest lasting 10 s was tested in multiple flies in the larval, early pupal, and late pupal stages (Supplementary Table 3). Excitation light power densities of 3.63 mW/mm^2^, 0.46 mW/mm^2^, and 7.26 mW/mm^2^, respectively were optimal for inducing continuous cardiac inhibition for 10 s.

**Supplementary Table 3.**
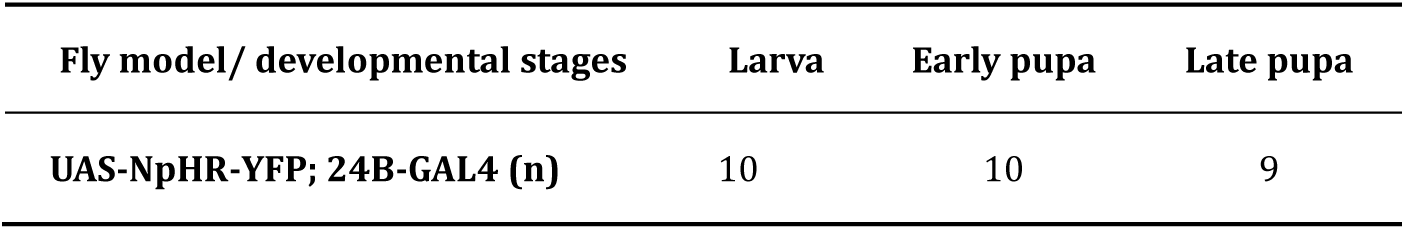
The number of UAS-NpHR-YFP; 24B-GAL4 flies measured in the larval, early pupal, and late pupal developmental stages for determining the needed excitation power density to induce cardiac arrest.

### IV. Supplementary video captions

A. **Supplementary video 1.mp4 –Red-light excitatory pacing of a ReaChR larval fly.** The heart of the larval fly beat at the resting heart rate (RHR) of 2.8 Hz initially, then followed the stimulation pulses at a frequency of 4.5 Hz, and returned to the RHR after the red-light stimulation was suspended.
B. **Supplementary video 2.mp4 –Red-light excitatory pacing of a ReaChR early pupal fly.** The heart of the early pupa beat at the RHR of 1.4 Hz initially, then followed the stimulation pulses at a frequency of 3 Hz, and returned to the RHR after the red-light stimulation was suspended.
C. **Supplementary video 3.mp4 - Red-light excitatory pacing of a ReaChR late pupal fly.** The heart of the late pupa beat at the RHR of 2.5 Hz initially, then followed the stimulation pulses at a frequency of 5 Hz, and returned to the RHR after the red-light stimulation was suspended.
D. **Supplementary video 4.mp4 - Red-light excitatory pacing of a ReaChR adult fly.** The heart of the adult fly beat at the RHR of 8.1 Hz initially, then followed the stimulation pulses at a frequency of 10.5 Hz, and returned to the RHR after the red-light stimulation was suspended.
E. **Supplementary video 5.mp4 - Red-light stimulation induces cardiac arrest in an NpHR larval fly.** The heart of the larval fly beat at the RHR of 4.8 Hz for the first 10 s with no red-light stimulation. It suspended beating immediately after the red light was turned on, and remained in a relaxed state for the 10s duration of red-light illumination. The heart resumed beating after the red light was turned off.
F. **Supplementary video 6.mp4 - Red-light stimulation induces cardiac arrest in an NpHR early pupal fly.** The heart of the early pupa beat at the RHR of 2.2 Hz for the first 10 s with no red-light stimulation. It suspended beating immediately after the heart was illuminated and remained in a relaxed state for the 10s duration of red-light illumination. The heart resumed beating after the red light was turned off.
G. **Supplementary video 7.mp4 - Red-light stimulation induces cardiac arrest in an NpHR late pupal fly.** The heart of the late pupal fly beat at the RHR of 1.6 Hz for the first 10 s. It suspended beating immediately after the heart was illuminated and stayed in a relaxed state for the 10s duration of red-light illumination. The heart resumed beating after the red light was turned off.
H. **Supplementary video 8.mp4 - Red-light inhibitory pacing of an NpHR larval fly.** The heart of the larval fly beat at the RHR of 3.4 Hz, slowed down to 1 Hz with a pulse frequency of 1 Hz and pulse duty cycle of 90%, and the heart rate returned to the RHR after the red-light illumination was suspended.
I. **Supplementary video 9.mp4 - Red-light inhibitory pacing of an NpHR early pupal fly.** The heart of an early pupa beat at the RHR of 2.7 Hz, slowed down to 1 Hz with a pacing frequency of 1 Hz and a pulse duty cycle of 80%, and the heart rate returned to the RHR after the red-light illumination was suspended.
J. **Supplementary video 10.mp4 - Red-light inhibitory pacing of an NpHR late pupal fly.** The heart of the late pupal fly beat at the RHR of 3.3 Hz, slowed down to 1 Hz with a pulse frequency of 1 Hz and a pulse duty cycle of 70%, and the heart rate returned to the RHR after the red-light illumination was suspended.
K. **Supplementary video 11.mp4 – Cardiac recovery after red-light stimulation for 1 s in an NpHR early pupal fly.** The fly heart rate quickly returned to the RHR after the red-light illumination was suspended.
L. **Supplementary video 12.mp4 – Cardiac recovery after red-light stimulation for 2 s in an NpHR early pupal fly.** The fly heart rate quickly returned to the RHR after the red-light illumination was suspended.
M. **Supplementary video 13.mp4 – Cardiac recovery after red-light stimulation for 5 s in an NpHR early pupal fly.** After the red-light illumination was suspended, an overshoot of HR was observed followed by a gradual HR recovery to the RHR.
N. **Supplementary video 14.mp4 – Cardiac recovery after red-light stimulation for 10 s in an NpHR early pupal fly.** After the red-light illumination was suspended, an overshoot of HR was observed followed by a gradual HR recovery to the RHR.
O. **Supplementary video 15.mp4 – Cardiac recovery after red-light stimulation for 20 s in an NpHR early pupal fly.** After the red-light illumination was suspended, an overshoot of HR was observed followed by a gradual HR recovery to the RHR.

